# Genetic surfing in human populations: from genes to genomes

**DOI:** 10.1101/055145

**Authors:** Stephan Peischl, Isabelle Dupanloup, Lars Bosshard, Laurent Excoffier

**Affiliations:** CMPG, Institute of Ecology and Evolution, Baltzerstrasse 6, 3012 Berne, Switzerland; Interfacultary Bioinformatic Units, Baltzerstrasse 6, 3012 Berne, Switzerland; Swiss Institute of Bioinformatics, 1015 Lausanne, Switzerland

**Author notes:** Correspondence Stephan Peischl & Laurent Excoffier Baltzerstrasse 6, 3012 Berne, Switzerland &.

**Keywords:** Founder effect, mutation load, expansion load, human expansions, human evolution

## Abstract

Genetic surfing describes the spatial spread and increase in frequency of variants that are not lost by genetic drift and serial migrant sampling during a range expansion. Genetic surfing does not modify the total number of derived alleles in a population or in an individual genome, but it leads to a loss of heterozygosity along the expansion axis, implying that derived alleles are more often in homozygous state. Genetic surfing also affects selected variants on the wave front, making them behave almost like neutral variants during the expansion. In agreement with theoretical predictions, human genomic data reveals an increase in recessive mutation load with distance from Africa, an expansion load likely to have developed during the expansion of human populations out of Africa.

## Main text

Since human populations have spread over the planet by a series of spatial range expansions, it is important to understand the consequences of these expansions on our genomic diversity. In a seminal paper, Cavalli-Sforza and colleagues [1] have shown that new mutations occurring on the edge of spatial expansion can increase in frequency and spread over a large portion of the newly colonized territories, far from their place of origin where they can even disappear. This increase in frequency of rare variants was later coined allele surfing [2], but we shall use here the more generic term of genetic surfing. It applies not only to new mutations, but to any standing variants happening to be on or close to the wave front of a range expansion [3–5], or to alleles introgressing from other species [6,7]. The genetic consequences of range expansions for single neutral loci are fairly well understood, and they have been described in several reviews [see e.g. 3,8–11]. In the present review, we will thus focus more on the causes and effects of surfing at the genomic level rather than at the single locus level, and contrast the surfing of neutral and selected variants, as range expansions can also affect the latter [12–15].

## Mechanism and consequences of genetic surfing

Genetic surfing results from the stochastic evolution of allele frequencies on expanding wave fronts where population densities are low and genetic drift is strong. It has been described in models of continuous habitat where population densities are monotonously declining towards the front of the expansion [4,5,14,16,17] (Fig. 1A), and in models with discrete habitats where the expansion proceeds by series of serial founder effects [18–23] (Fig. 1B). Note that from a modelling perspective, genetic surfing is conceptually similar to the hitchhiking of neutral (or deleterious) alleles with beneficial variants that sweep through a spatially structured population [24,25]. Due to the sampling of individuals colonizing new territories and high rates of genetic drift on the front, some derived alleles are lost at some loci, and some alleles are increasing in frequencies at others. It is this increase in frequency over time and space at those loci where derived alleles are not lost that has been called genetic surfing. It is difficult to define genetic surfing more precisely than this propagation of alleles conditional on non-loss [19], as some alleles can spread over areas of variable size and reach variable frequencies [2]. In extreme cases, derived alleles can fix on the wave front and surf over very large distances, leading to sectors of no or very low diversity in two dimensional expansions [4,5] (Fig. 1C). Range expansions thus globally lead to a decrease of genetic diversity along the expansion axis [26] due to the stochastic loss of variants. Genetic surfing is favored by low effective sizes on the front, low migration rate from the back of the wave, and high levels of growth on the wave front [2,19,27], but Allee effects and long distance dispersal can limit or prevent surfing and preserve diversity [28–34].

**Figure 1.**
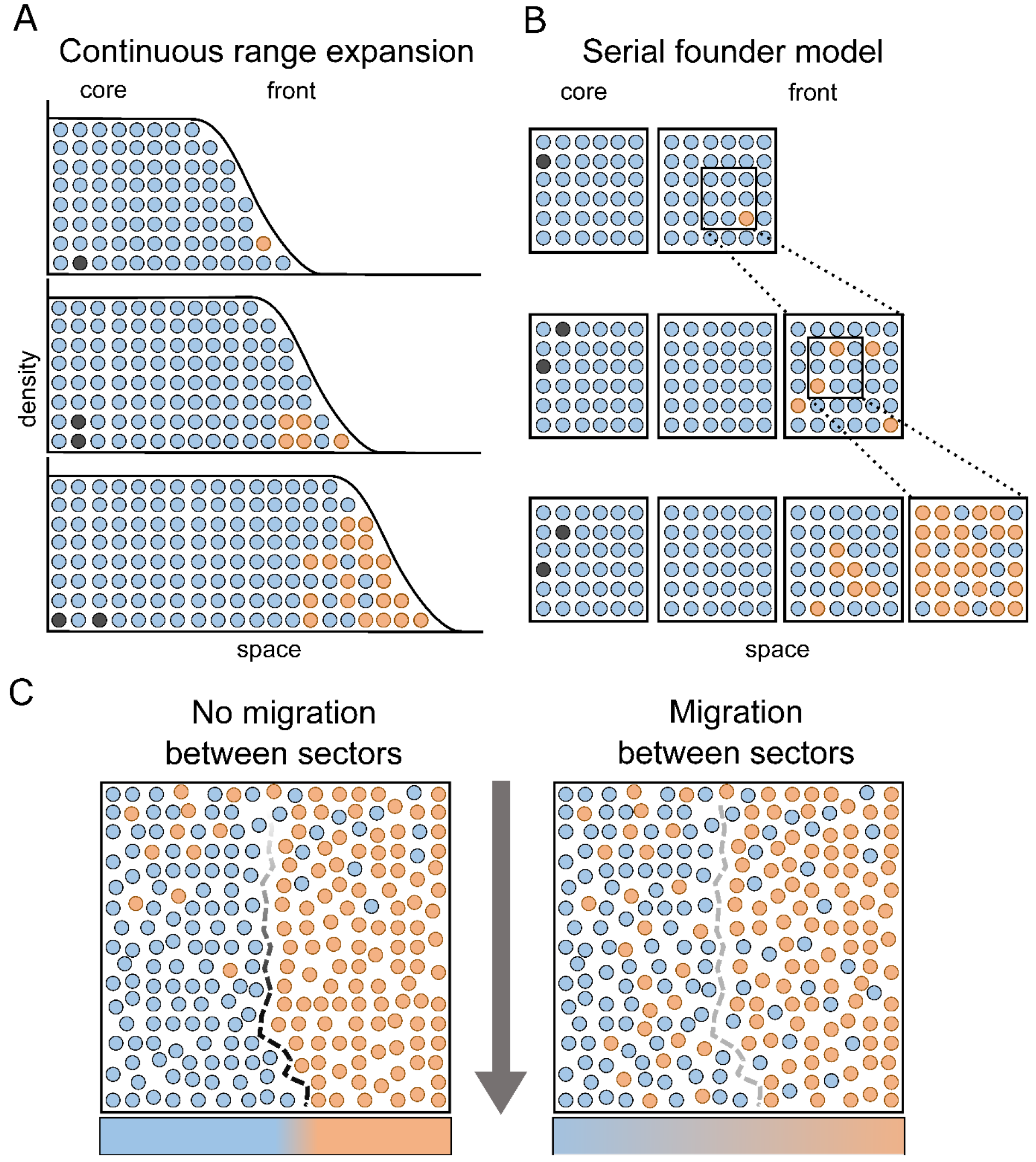
Allele frequency changes due to genetic surfing.**A**: During a range expansion in a continuous one dimensional habitat, a new mutation appearing onthe wave front (orange) can increase in frequency and surf on the wave of advance, reaching high frequencies in newly occupied territories. **B**: Discrete serial founder model of range expansion. A given number of founders are sampled from the leading deme to colonize an empty deme on the right. These founders then reproduce, and a new set of founders is selected to colonize the next deme. Note that in both models A and B, a new mutation appearing in the core of the range (black) has a much lower probability to increase in frequency than a new mutation in the front (orange), because individuals in the core have less descendants than those on the front. **C**: Two-dimensional expansions from top to bottom, starting from a well-mixed population. Without migration between colonized areas (left pane),the expansion leads to the (local) fixation of a random neutral allele by surfing and thus tothe creation of sectors of low diversity. The sectors are delineated by a dashed line whose shade of grey indicates the intensity of the differentiation between sectors. With lateral migrations between colonized areas, the sectors do not emerge as clearly, and a transitory gradient of diversity perpendicular to the expansion axis can establish (bottom row).

## Surfing of standing vs. new mutations

At neutral loci, the overall frequency of derived alleles in the genome that are already present at the onset of a range expansion (standing variants) should not change under the sole effect of drift, founder effects, or bottlenecks because the loss of derived alleles at some loci is compensated by the increase in frequency of derived alleles at other loci. Individuals on the front of an expansion should therefore have a smaller number of polymorphic sites than individuals from the core, but they should be more often homozygote for derived alleles. Overall, they should thus have similar numbers of derived alleles in their genome than individuals in the core (Fig. 2). Front individuals should also have higher frequencies of derived alleles at sites shared with core individuals, a property that has been exploited to polarize the direction of range expansions and to locate their geographic origin [35]. Individuals have a higher chance to transmit their genes to later generations if they are present on a wave front, because they have a higher reproductive success than individuals from the core [36]. It may therefore appear as if genetic surfing is a non-neutral process because any given allele that finds itself on the wave front experiences an immediate increase in absolute frequency, due to the higher growth rate on the expansion front than in the core. However, this apparent increase in absolute fitness is only visible if we condition mutations to occur in founding individuals, and vanishes if we take into account that many mutations at the wave front are not sampled during later founder events (Figs. 1A and Figs. 1B, see also s #x003D; 0 in Fig. 3).

**Figure 2.**
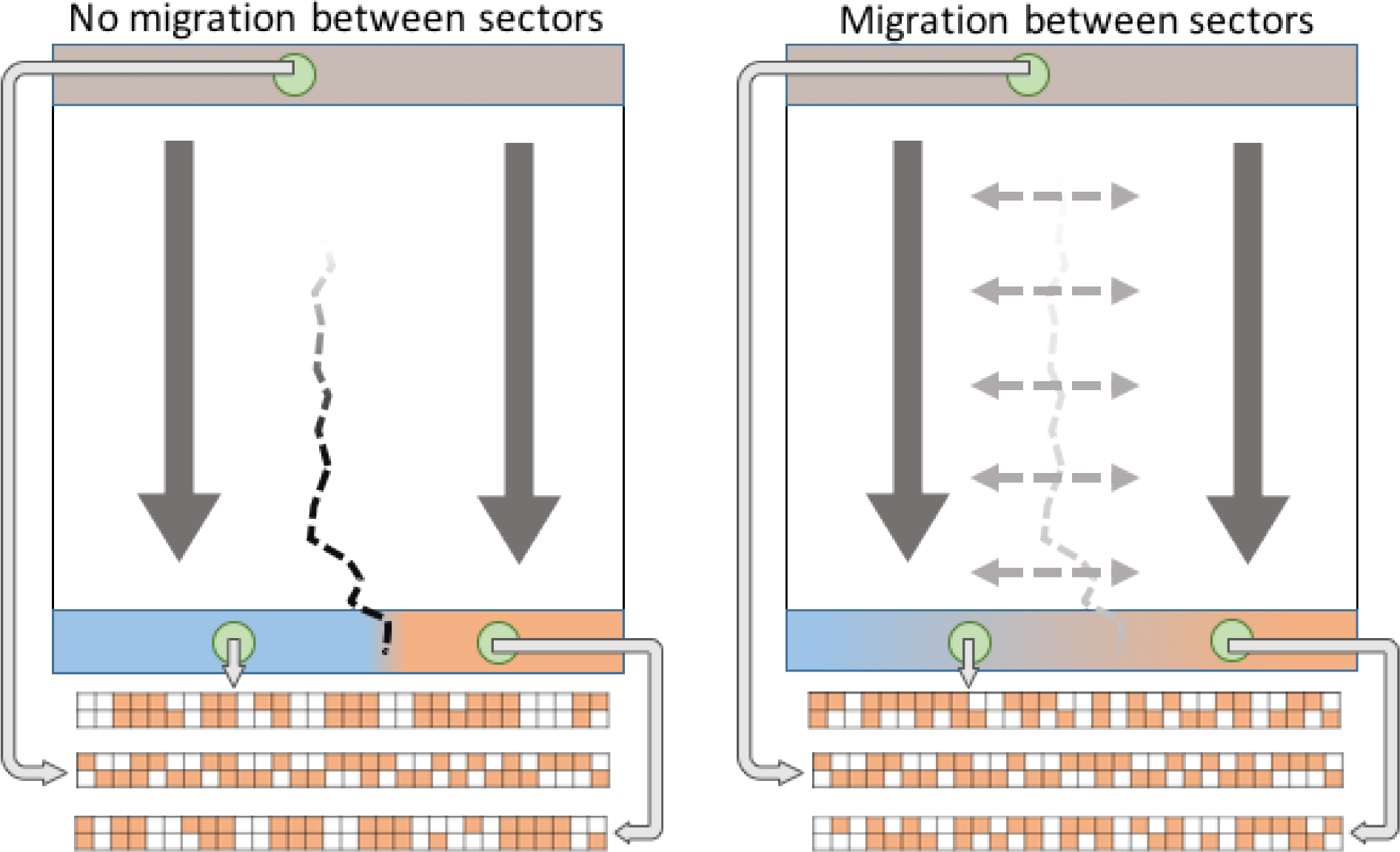
Illustration of the type of genomic diversity expected in core and front individuals after surfing under scenarios represented in Fig. 1C. Vertical arrows represent the direction of the expansion, whereas dashed horizontal arrows in the right pane represent lateral gene flow between neighboring demes. We show here the putative genomic diversity of diploid individuals genotyped at 30 loci either sampled from the core (top green circle) or from the front (bottom green circles). Derived alleles are denoted as orange squares. The expected total number of derived alleles is the same in all individuals, but due to surfing, front individuals have less sites with derived alleles, and thus more loci with homozygous derived alleles than typical core individuals. Derived allelefrequencies at sites shared between front and core populations are therefore larger in front than in core populations [19]. In presence of migrations between sectors (right pane), the number of polymorphic sites on the front is higher than without lateral gene flow, and heterozygosity is restored more rapidly.

## Gradients of diversity due to surfing

The fact that genetic diversity should decrease by allele loss almost linearly with distance from the origin of an expansion [18,20] has been used to identify the origin of human expansions out of Africa. Early studies of microsatellite diversity have revealed an almost perfect linear decrease of heterozygosity with distance from Ethiopia [21,37], which has been confirmed with genomic data[e.g.38]. Interestingly, human range expansions are also associated with gradients of linkage disequilibrium [39,40], recombinational diversity [41], length of runs of homozygosity (ROHs) [42], allele frequencies [35,43], as well as a right shift or a flattening of the site frequency spectrum [38,44]. Given these multiple evidence for the creation of gradients of diversity along expansion axes, one would expect that the largest extent of differentiation should be between the source and the edge of an expansion. However, PCA analyses of simulated data reveal that the main axis of differentiation after an expansion runs perpendicular to the expansion axis [45], which is explained by the differentiation existing between surfing sectors along the expansion axis (see Fig. 1C). Consequently, the classical PC analyses of human European diversity showing gradients of differentiation from the Near-East to North Western Europe [46] are unlikely to be the signature of a Neolithic demic diffusion [45], but could rather reflect re-expansions (e.g. post-LGM) from different refuge areas in Southern Europe [47].

## Surfing or adaptive evolution?

A striking feature of genome scans for positive selection in humans is that they reveal many candidate SNPs that have very similar geographic distributions, with high contrasts observed between continental groups [48,49], and where Africa and America show the largest number of loci with extreme frequency differences [50]. Simulations have shown that such large continental differences could simply arise from surfing of neutral variants during the out of Africa expansion [51]. Therefore, previous claims of signals of adaptation based mainly on the spatial distribution of allele frequencies [e.g. 52,53] should probably be taken with great caution or revised [54], and could partly explain some lack of consistency between the outcome of various genome scans [55]. It has also been recently recognized that there was an overall lack of evidence for strong signals of selective sweeps in the human genome [56–58], with only a handful of regions with fixed differences between continents [59]. It is difficult to understand why positive selection, if active, has not led to more fixations of beneficial mutations in human populations even under polygenic selection models [60]. A low number of fixed difference due to genetic surfing between the source of human expansions (Africa) and the edges of expansions is equally puzzling, as one would expect that surfing mutations should quickly fix on wave fronts (Fig. 3) [1,2,19]. However, mutations can surf in large populations without necessarily being fixed on the front [2], and even if different mutations fix in different sectors, migration between parallel sectors can lead to intermediate final frequencies in newly colonized areas (Fig. 1C). In addition, some long distance dispersal events between demes in the wake of the expansion might have reshuffled diversity and limited the fixation of derived alleles in non-African populations [61].

**Figure 3.**
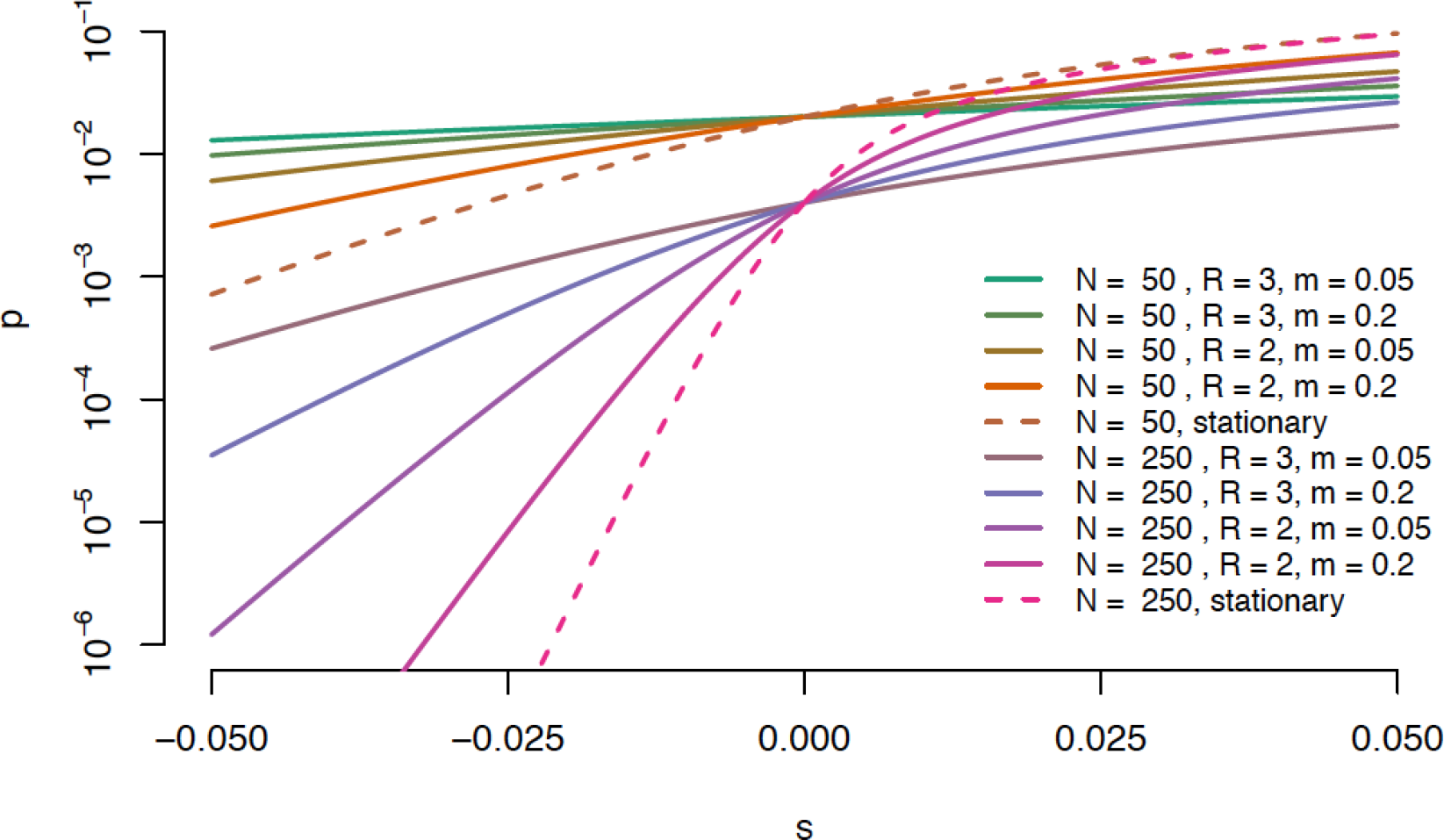
Probability of fixation on the wave front for a new mutation occurring at the wave front during a one dimensional range expansion (serial founder model) for deleterious, neutral and advantageous variants (figure based on equation (1) in ref [15]). Mutations are assumed to occur in a fully colonized deme just before the sampling of founders such that they have an initial frequency of 1/2N (Fig 1B).The dashed line represents the probability of fixation in a single stationary deme, which is smaller for deleterious mutations and larger for beneficial ones.

## Surfing of deleterious mutations leads to an expansion load

Genetic surfing is not restricted to neutral variants. The probability that a mutation establishes itself on the expansion front and spreads with the expanding wave is largely driven by genetic drift, which makes the rate at which beneficial, neutral or deleterious mutations are established at the expansion front more similar to each other relative to what is expected in the core [15] (Fig. 3). Mildly positively or negatively selected variants therefore mostly behave like neutral variants on the front of the expansion. Deleterious variants can therefore temporarily surf on the expansion wave due to the high growth rate prevailing in front populations [12]. In asexual (non-recombining) organisms with high mutation rates, genetic surfing can even lead to an accumulation of deleterious mutations at the expansion front [14]. Interestingly, and perhaps somewhat conter-intuitively, beneficial mutations that occur on the front of an expansion have a higher probability to become permanently established in the whole population (even if they fail to surf) as compared to mutations that occur in the core [13]. Indeed, beneficial mutations occuring on the wave front quickly increase in absolute frequency due to high growth rates in these marginal populations (Fig. 1A, 1B), which lowers the probaility that they are initially lost. This effect is similar to what has been observed in demographically growing well-mixed populations [62]. The spread of beneficial mutations in the core has, however, little impact on the evolution at the expansion front because beneficial mutations sweep through the core at a speed [25] that is much lower than that of the expanding wave [14,63,64]. Since beneficial mutations that establish on the front spread much faster than beneficial mutations in the core [14,65], they provide opportunity for rapid adaptation during early stages of an expansion [66]. In the long run, however, genetic drift at the expansion front dominates and, since beneficial mutations occur at a lower rate than deleterious ones [67], the overall fitness of the populations should decrease along the expansion axis under a broad range of conditions [15]. The resulting elevated mutational burden of expanding species has been coined expansion load [15]. Expansion load generally builds up fastest under conditions that favor genetic surfing, such as small local carrying capacities, high growth rates, and low migration rates [2], which contributes to making the probabilities with which deleterious and beneficial alleles fix on the wave front relatively similar to that of neutral alleles (Fig. 3).

In the absence of epistasis or dominance, only new mutations contribute to expansion load [63], and the resulting decrease in fitness is relatively slow. If some deleterious mutations are (partially) recessive [63], the increase in homozygosity of standing deleterious variants along the expansion axis leads to a very rapid increase of the (recessive) mutation load [63]. Another interesting consequence of recessivity is the establishment of heterozygosity-fitness correlations during 2D expansions: migration between sectors in which different mutations have established (see Fig. 1C and Fig 2) creates a ridge of high fitness #x201C;hybrids#x201D; in which the effects of deleterious mutations are partially masked [63]. Whereas this effect has been oberved in plants after a range expansion [68], it remains to be shown in humans.

Even though expansion load is a transient pheomenon, simulations show that the increased mutation load can persist for hundreds to thousands of generations after the expansion has ended [15]. The rate of purging of deleterious mutations depends on factors such as selection coefficients, the degree of dominance and the amount of gene flow between populations [15,63,69–71].

**Figure 4.**
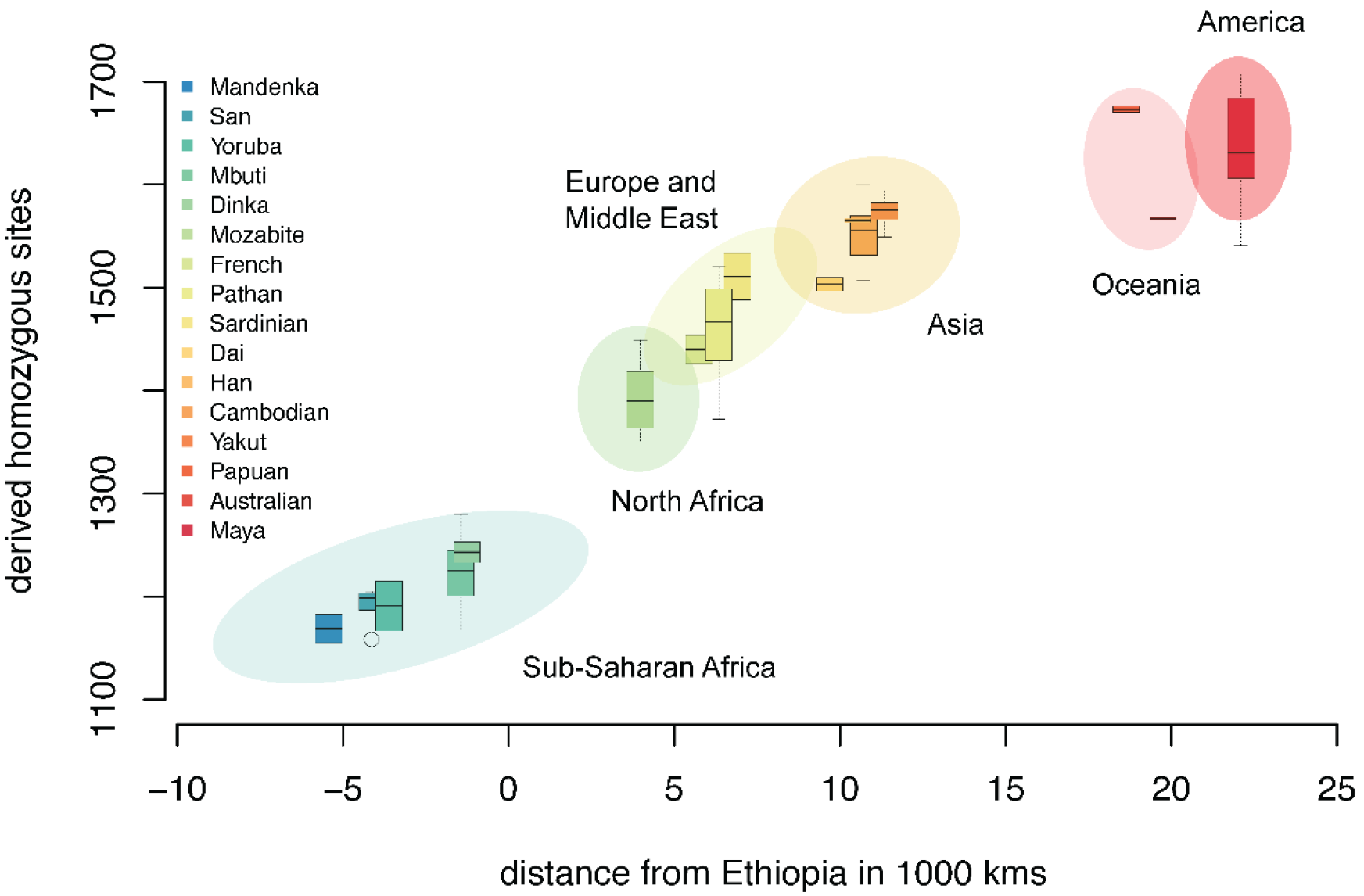
Recessive mutation load as a function of the least-cost distance from Addis-Ababa (Ethiopia). Negative distances denote sub-Saharan-African populations. Recessive mutation load is measured as the number of sites homozygote for derived mutation at sites that have a GERP rejected substitution score [72] larger than 4, indicative of relatively strong evolutionary constraints. Reported populations are from [44], complemented by samples from [73,74], all individuals examined for the same 44Mb of exomic sequences without missing data (see [44] for details).

## Evidence for expansion load in human populations

Evidence for expansion load is generally difficult to get as fitness is difficult to measure directly. Ideally, one would need to evidence the spread over a wide area of attested deleterious mutations in populations with a history of range expansions. In Europe, a couple of disease mutations are increasing in frequencies along a South-East to North-West gradient (e.g. *HFE-C282Y* associated to hematochromatosis [75] or ΔF508 associated to cystic fibrosis [76]), and a CPT1A mutation associated with hypoglycemia and high infant mortality has been shown recently to have greatly increased in frequency in arctic populations [77]. Interestingly, in the three cases mentioned above, the spread of these deleterious variants over long distances has been explained by past episodes of positive selection, but genetic surfing of only deleterious variants seems an equally reasonable and more parsimious hypothesis.

Another way to approximate mutation load from sequence data is to estimate the degree of evolutionary conservation of mutations at particular positions in the genome (see e.g., ref [78]). Using such an approach, a clear gradient of homozygosity at predicted deleterious sites has been observed along the human expansion axis out of Africa [44] (Fig. 4), in keeping with predictions [63]. Furthermore, this cline becomes weaker with increasing deleteriousness of mutations [44], suggesting that the evolution of mildly and moderatly deleterious mutations are dominated by strong genetic drift during the expansion, whereas selection still dominates the evolution of strongly deleterious muations, as predicted by theory [15].

The impact of demographic history on mutation load is still a hotly debated topic [44,71,78]. When focusing on the number of predicted deleterious alleles per individuals, which is equivalent to considering that load is due to co-dominant mutations, no differences were found between Europeans, Americans of European ancestry or African-Americans [80,81,84]. However, the relative proportion of alleles predicted deleterious is significantly higher in Europeans as compared to Sub-saharan Africans [15,79]. These apparently contradictory results can be understood from Fig. 2, by realizing that while each individual carries roughly similar numbers of derived alleles before and after surfing, these alleles are distributed over fewer loci in populations having gone through a range expansion (e.g. Europeans) as compared to core populations (e.g. Sub-Saharan Africans), and therefore reach higher frequencies in front populations [19]. Deleterious alleles that are kept at very low frequency in core populations and that have increased in frequency after surfing have a higher probability of being detected in a sample of individuals on the front, contributing to the observation of a larger proportion of non-synonymous and potentially deleterious alleles in non-African populations [15,38,79].

The total genetic load of an individual and its fitness will also highly depend on the distribution of dominance coefficients in the genome [44,71]. If all mutations were codominant, one would expect no or little differences in genetic load between populations, because mildly deleterious mutations have mainly evolved as neutral mutations during theand expansion (or during a bottleneck [81]). However, the mutation load should be clearly larger in front populations if some of the surfing deleterious mutations are partially recessive. The extent of this load excess in front populations is difficult to assess since recessivity appears to be correlated with deleteriousness [85]. The most deleterious variants, which should be the most recessive ones, should not be influenced much by surfing since their frequencies are maintained consistantly low in all populations by selection [44]. The total genomic frequency of very midly deleterious variants and their effect on load should also not vary due to surfing as they should be almost co-dominant. It thus likely that difference in load between populations should be mainly due to partially recessive variants of intermediate effects (e.g. [71]). These predictions offer interesting areas of research in pheno-genomic diversity, showing the need of better methods to asess the phenotpic effect of mutationsmutation in humans, and to distinguish between the effects of somatic and germ-line mutations [86].

## Acknowledgements

ID and LB were partially supported by a Swiss NSF grant No 31003A-143393 to LE.

